# Transcriptomic analysis of adult mouse cardiac stromal cells using single-cell qRT-PCR

**DOI:** 10.1101/2024.12.22.629898

**Authors:** Rita Alonaizan, Patricia Chaves-Guerrero, Sara Samari, Michela Noseda, Nicola Smart, Carolyn Carr

## Abstract

Fate mapping studies have challenged the longstanding view of the adult mammalian heart as a post-mitotic organ, suggesting limited cardiomyocyte renewal. This has spurred efforts to identify cardiac progenitor cell (CPC) populations, but their contribution to cardiac regeneration has been found to be minimal compared to cardiomyocyte proliferation. Despite this, CPC transplantation has shown therapeutic potential through paracrine signalling.

The identity of CPCs remains unclear due to overlapping characteristics with other cardiac stromal cell populations such as fibroblasts, mesenchymal cells, and pericytes. This study sought to optimise the isolation of CPCs by developing a cardiac collagenase-trypsin (CT) protocol, which was compared to the established method of isolating cardiosphere-derived cells (CDCs). The CT protocol resulted in a higher cell yield and reduced expansion time, with both CTs and CDCs showing superior survival potential under serum starvation compared to commercially acquired cardiac fibroblasts (CFs).

Single-cell qRT-PCR analysis revealed that CTs and CDCs share a similar gene expression profile, distinct from CFs, characterised by the enrichment of cardiogenic transcription factors. Notably, CTs exhibited higher expression of *Tcf21* and lower *Tbx5*, suggesting an epicardial-derived fibroblast phenotype, whereas *Tbx5* was enriched in CDCs and CFs. Additionally, CTs showed an enrichment of macrophage-associated genes *Mrc1* and *Csf1r*, possibly due to the transdifferentiation of macrophages to or from a fibroblast phenotype in a subset of CTs. The study concludes that CTs represent a robust and efficient source of CPCs with therapeutic potential and offers insights into the complex identity of cardiac stromal cells.

## Introduction

Evidence stemming from fate mapping studies of cardiomyocyte renewal in the adult mammalian heart challenged the dogma considering the heart as a post-mitotic organ^1–3^. Therefore, multiple research groups attempted to characterise endogenous cardiac populations that may have a role in the regenerative process. Identifiers of the presumed cardiac progenitor cell (CPC) populations include SCA1 and the SP dye-efflux phenotype, KIT, ISL1, cardiosphere- and colony-forming assays, or re-activation of the embryonic epicardial programme and WT1 upregulation^4^. More recent findings indicate that the mechanism behind cardiomyocyte renewal is cardiomyocyte proliferation, and that CPC contribution is minimal^1–6^. Despite their low cardiomyogenic potential after transplantation into the infarcted heart, CPCs have demonstrated therapeutic benefits in pre-clinical studies, primarily due to paracrine signalling^7–11^.

The identity of CPCs remains unclear, with numerous studies questioning the distinctions between CPCs and various cardiac stromal cell populations, including mesenchymal cells, fibroblasts, and pericytes^12–16^. This uncertainty stems from the overlap in their morphology and gene expression profiles. For instance, cardiac fibroblasts, like CPCs, express cardiogenic transcription factors that contribute to cardiac development and repair^17^. Both cardiac and tail fibroblasts share a molecular signature similar to that of mesenchymal stem cells (MSCs)^18^. Significant efforts have been made to distinguish these cardiac stromal cells. To complicate matters further, single-cell transcriptional profiling of mouse ventricular non-myocytes has revealed subpopulation heterogeneity within the cardiac fibroblast population^19^. Additionally, markers commonly used to identify pericytes are relatively non-specific, albeit some genes are enriched in pericytes compared to other cell types^19^.

Certain research groups have sorted CPCs based on surface marker expression followed by clonogenic selection, which include the derivation of SCA1+ PDGFRα+/SP cells^9,20^. These methods produced homogeneous, self-renewing cardiogenic populations that showed promise. However, these cell types are found at very low rates in the adult mouse heart, with a reported cloning efficiency of around 1-2%. Ideally, an optimal cell isolation protocol would yield a large, robust, and reproducible population of cells in the shortest possible duration. The isolation of cardiosphere-derived cells (CDCs) involves culturing cardiac explants to harvest explant-derived cells (EDCs), which are then induced to form cardiospheres. EDCs vary in morphology and time required to achieve confluent culture, with CDC expansion times ranging from 5 to 84 days^21^.

To optimise CPC isolation, we developed a cardiac collagenase-trypsin (CT) protocol adapted from studies by Gharaibeh *et al*.^22^ and Okada *et al*.^23^ for isolating slowly adhering cells (SACs) from skeletal muscle. These SACs demonstrated superior differentiation, survival, and therapeutic potential compared to rapidly adhering cells (RACs). Comparably, we showed that these CPCs, which we term CTs, can be induced to differentiate into the cardiomyocyte lineage *in vitro*^24^ using TGFβ1 stimulation^25,26^. We also showed that fatty acid supplementation triggered a metabolic switch via the PPARα pathway, evidenced by upregulated oxidative metabolism and increased expression of cardiomyocyte markers in CTs^24^. This aligns with the metabolic switch from glycolysis to fatty acid oxidation observed during cardiomyocyte maturation in development^27^. In this study, we characterise the presumed CPC populations, CTs and CDCs, in comparison to commercially acquired primary cardiac fibroblasts (CFs) (**Figure 1**) using single-cell qRT-PCR and cell survival assays. Our aim is to profile CPCs obtained without clonogenic and marker selection and compare them to the cardiac fibroblast compartment.

**Figure 1.**
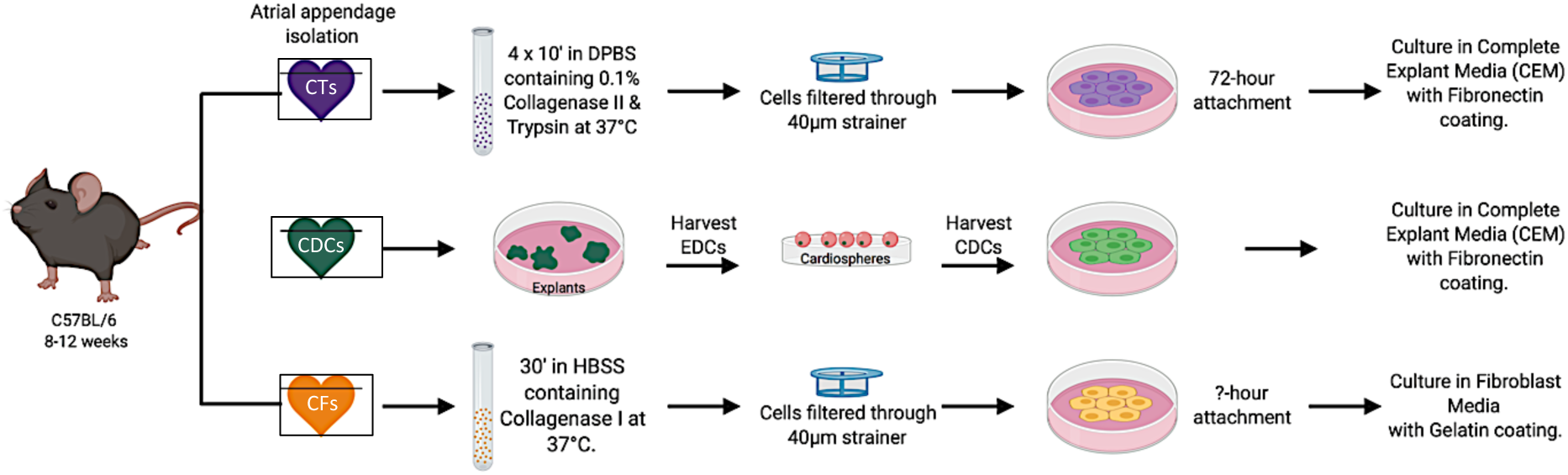
Isolation method of the three cardiac stromal populations, CTs, CDCs and CFs. Schematic showing the protocol used to isolate CTs, CDCs, and CFs. The isolation protocol of CTs and CDCs is detailed in the Methods. As CFs were purchased and not isolated in our lab, some information is lacking e.g. age of mice, duration of cell attachment, and composition of the accompanying media. Illustration created with BioRender.com.

## Results

### The CT protocol produces higher cell yield compared to CDCs, but a similar survival potential

CTs and CDCs were isolated from adult mouse atrial tissue and expanded for further analysis. The average time to reach 90% confluency at passage 0 and passage 1 was considerably shorter for CTs, taking approximately 10 days and 20 days, respectively. In contrast, CDCs required around 40 days to passage 0 and 60 days to passage 1, due to the additional steps involved in explant culture and cardiosphere formation (**Figure 2.A**).

**Figure 2.**
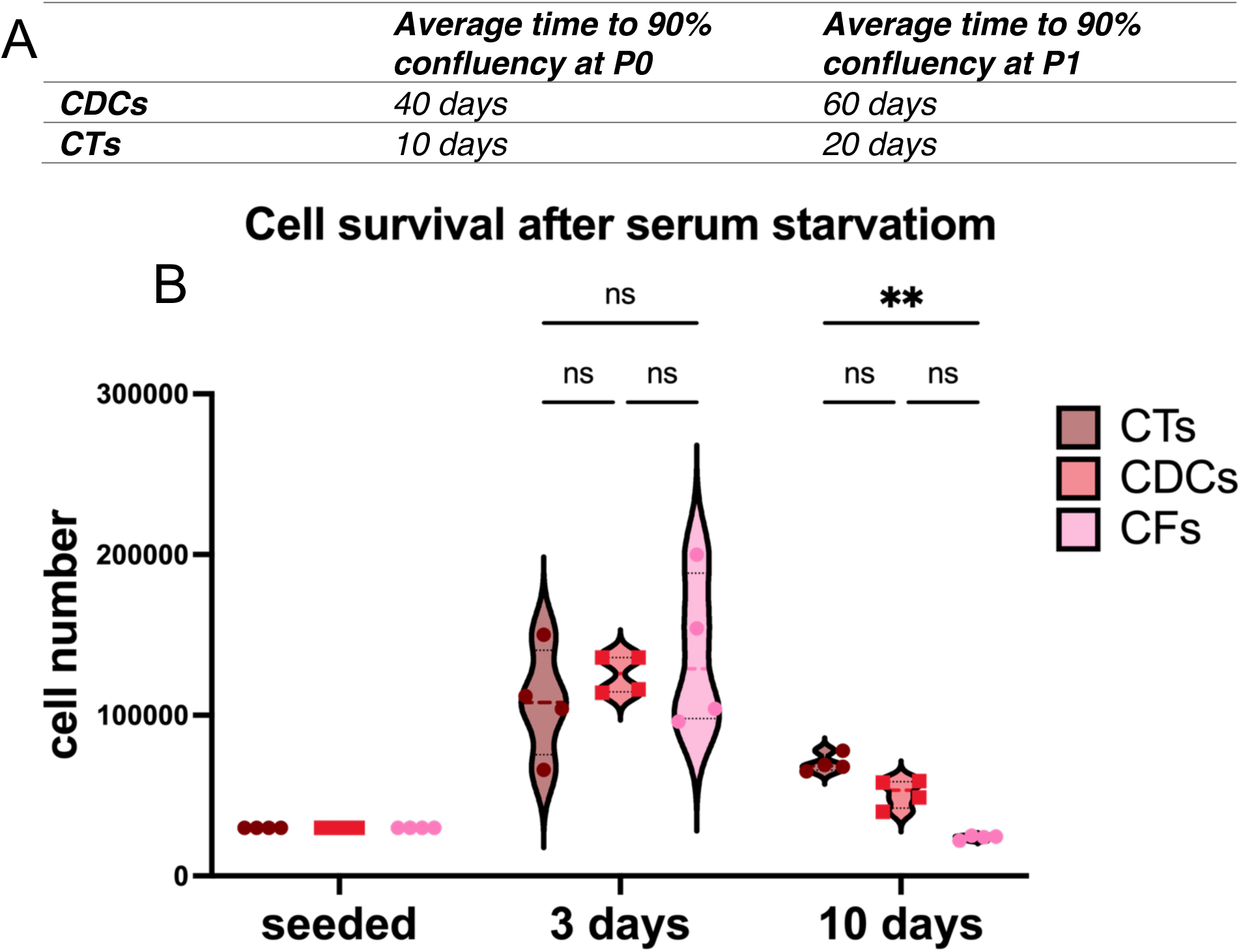
The CT protocol produces higher cell yield compared to CDCs but a similar survival potential. **(A)** Average time to reach 90% at passage 0 and passage 1, as assessed by cell counting, in CTs and CDCs. **(B)** Cell Survival following serum starvation for 3 or 10 days, as assessed by cell counting, in CTs, CDCs and CFs. Analysed using a two-way ANOVA with Tukey’s multiple comparisons test to determine statistical significance (n=4), * p < 0.05, ** p < 0.01.

After expansion to passage 3, we assessed the survival potential of CTs compared to CDCs and commercially acquired CFs to determine their robustness for transplantation. Serum starvation was used to challenge the cells in culture for 3 days and 10 days. CTs, CDCs, and CFs continued to grow following serum starvation with no significant difference in cell numbers detected at day 3. At day 10, CT cell numbers were significantly higher than CFs but not CDCs, suggesting that CTs have a superior survival potential compared to CFs (**Figure 2.B**).

### Single-cell profiling of cardiac stromal cells

CTs, CDCs and CFs isolated from adult mouse atrial tissue were expanded to passage 3 for single-cell qRT-PCR analysis and compared with the HL-1 cardiomyocyte line which was used as a control (**Figure 3**). Genes selected for this experiment (detailed in supplementary table 1) included those encoding cardiogenic transcription factors, stem cell-related markers, fibroblasts, cardiomyocytes and other genes that were differentially expressed in the SCA1 subpopulations according to the Noseda *et al*. study^9^. Moreover, we attempted to detect different cardiac subpopulations including pericytes, endothelial cells and macrophages within the cardiac stromal cell populations. The heatmap (**Figure 3.A**) shows that the HL-1 cardiomyocytes were distinct from the cardiac stromal cells with expression of each gene across the different populations shown in **Figure 3.B**. The cardiac stromal populations expressed the fibroblast-associated genes *Cd44*, *Ddr2*, *Pdgfrb*, *Acta2* (encodes αSMA), and *Vim*, albeit CFs had less consistent expression of *Pdgfrb*, and *Acta2*. Interestingly, *Ddr2, Acta2* and *Vim* were also expressed by HL-1 cardiomyocytes. Additionally, on rare occasions, CFs (n=2/29) and CTs (n=3/55) co-expressing *Wt1* and *Tcf21* were detected, perhaps inferring an epicardial-derived fibroblast identity. *Ly6a* (encodes SCA1) and *Pdgfra* were expressed in all three populations. The three cardiac stromal populations expressed little or no *Prg4* and *Wif1* but displayed expression of *Medag* and *Col1a1*. *Ms4a4d* expression was detected in CTs and some CFs only. Regarding pericyte-associated markers, *Kcnj8* showed little or no expression in all populations, *Colec11* showed sporadic expression, whereas *Ng2* showed high expression in all three cardiac stromal populations.

**Figure 3.**
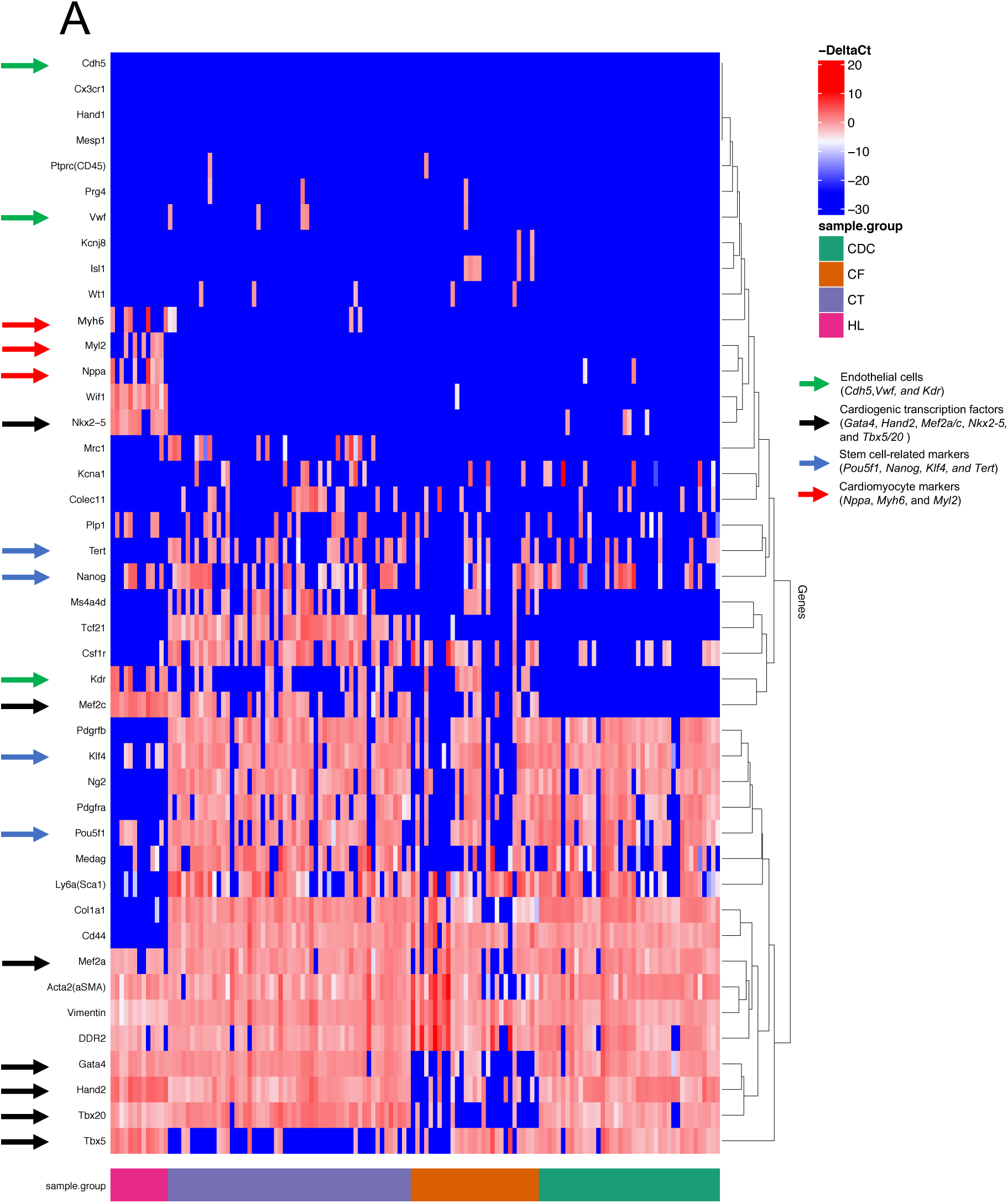
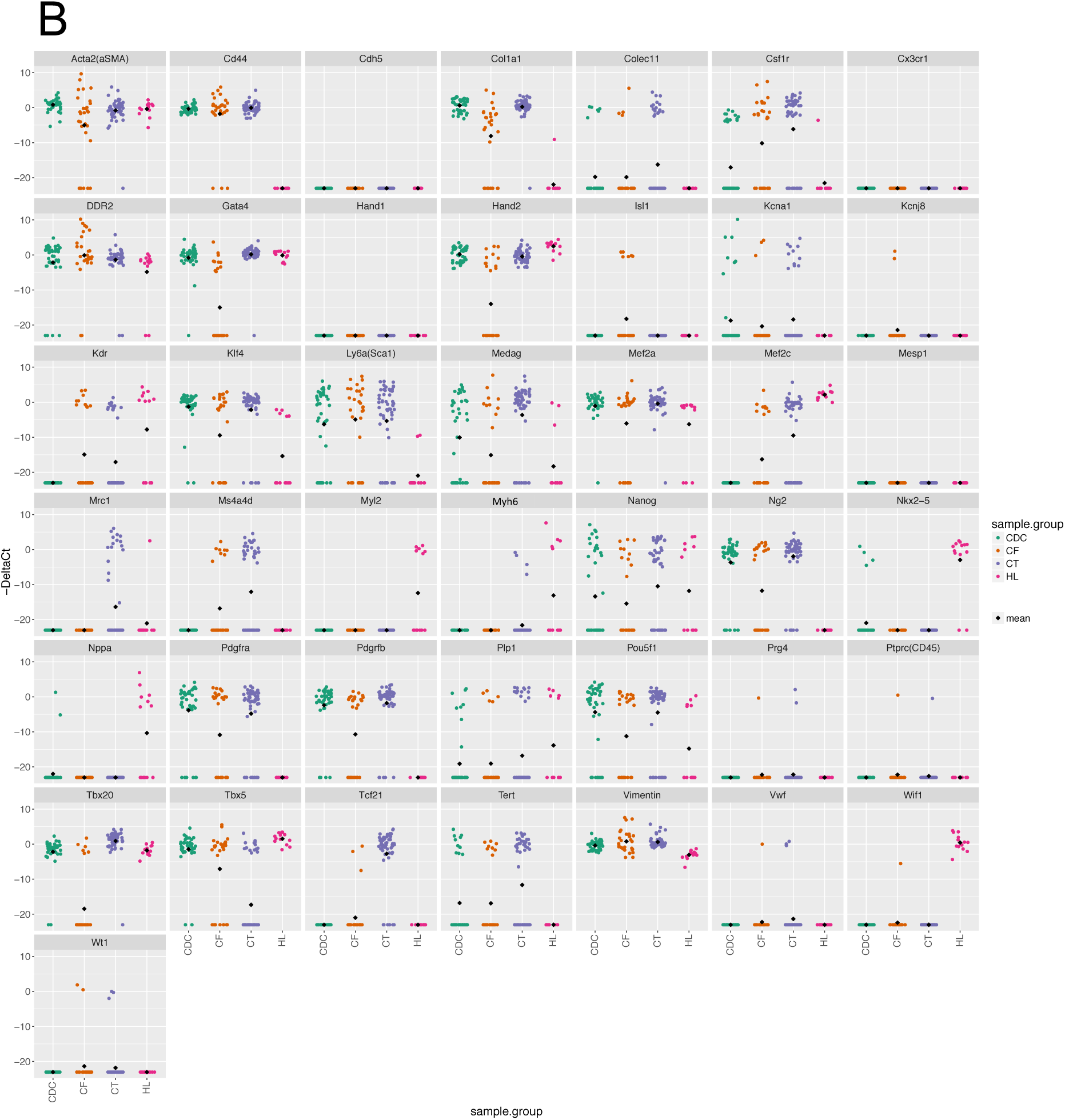
Single-cell analysis reveals an enrichment of Tcf21 and low expression of Tbx5 in CTs, compared to CDCs and CFs. **(A)** Expression of 43 genes was analysed at the single-cell level in the four cell populations shown: CTs (n=55), CDCs (n=41) and CFs (n=29) at passage 3, in addition to HL-1 cardiomyocytes (n=13). Both CTs and CDCs represent 3 biological replicates. The heatmap illustrates expression as -ΔCT values (blue is low or absent; red is high). Samples were ordered based on cell type, and the genes grouped by a hierarchical clustering algorithm based on the underlying co-expression pattern. **(B)** Single-cell expression of the 43 genes tested, represented individually in the four cell populations shown with the mean represented as a black diamond in each set.

In CTs and CDCs, the most prevalent cardiogenic transcription factors were *Gata4*, *Tbx20*, *Hand2*, and *Mef2a* with little or no expression of *Hand1*, *Isl1*, and *Nkx2-5*. In contrast, CFs had less uniform expression of *Gata4*, *Tbx20*, and *Hand2*. Unlike CDCs, both CTs and CFs expressed *Mef2c* heterogeneously. The expression of *Tbx5* in CFs followed a similar pattern to CDCs but was more sporadic in CTs. Interestingly, only CFs expressed *Isl1,* which has a very limited expression profile in the adult heart^28,29^. The three cardiac stromal populations had very little or no expression of markers of cardiomyocytes (*Nppa*, *Myh6*, and *Myl2*), haematopoietic (*Cd45*) and endothelial cells (*Cdh5* and *Vwf*), except for *Kdr* expression, which was detected in the CTs, CFs and HL-1 cardiomyocytes. Expression of stem cell-related markers (*Pou5f1*, *Nanog*, *Klf4*, and *Tert*) was found in the three cardiac stromal populations. *Pou5f1*, *Nanog*, and *Klf4*, but not *Tert*, were also expressed in HL-1 cardiomyocytes. As expected, HL-1 cardiomyocytes had a distinctly different expression profile and displayed expression of most cardiogenic transcription factors (*Gata4*, *Hand2*, *Mef2a/c*, *Nkx2-5* and *Tbx5/20*), and cardiomyocyte markers (*Nppa*, *Myh6*, and *Myl2*).

Expression of Schwann cell-related markers (*Plp1* and *Kcna1*) was found sporadically in all populations, except for HL-1 cardiomyocytes where only *Plp1*, but not *Kcna1*, was expressed. We also assessed the expression of the macrophage-associated genes *Mrc1*, *Csf1r* and *Cx3cr1*. *Cx3cr1* showed no expression in any population, and *Mrc1* was only expressed in CTs. In contrast, *Csf1r* was expressed in all three stromal populations but showed a more uniform pattern in CTs.

### A comparison of the molecular profiles of CPCs and CFs

By principal component analysis (PCA), cardiac stromal cells and HL-1 cardiomyocytes were resolved as discrete groups, which is consistent with their distinct phenotypes. Gene loadings contributing to each dimension (Dim) suggest that a small subset of genes explain the cross-group variability captured by Dim.1 and Dim.2. The separation of HL-1 cardiomyocytes was attributable to *Wif1*, *Nkx2-5*, *Mef2c*, *Nppa*, *Myl2*, and *Myh6* (**Figure 4.A**). Some separation of CFs from CTs was also revealed, with the CDCs clustering between the other two populations. This can be seen more clearly in the PCA of the stromal populations alone (**Figure 4.B**). The separation of CTs was attributable to *Tcf21*, clearly seen in **Figure 4.B**, while the separation of CDCs and CFs was mainly attributable to *Tbx5*. This *Tcf21* versus *Tbx5* polarity may reflect functional or origin heterogeneity. CFs appeared more dispersed, suggesting higher heterogeneity, in comparison to CTs and CDCs, which again is more apparent in the PCA of stromal populations only.

**Figure 4.**
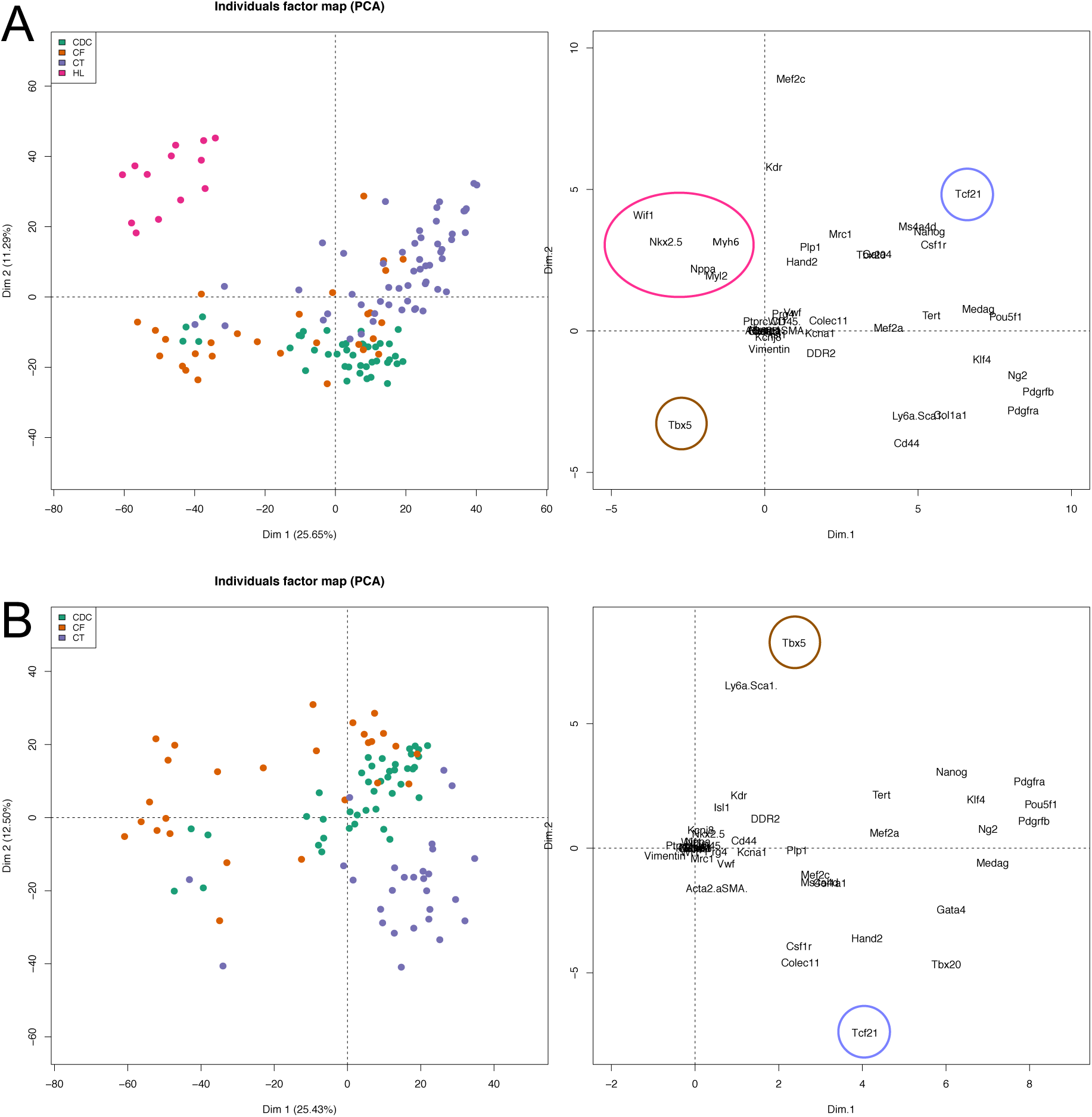
PCA of the three cardiac stromal populations and HL-1 cardiomyocytes. **(A)** PCA of the four cell populations shown: CTs (n=55), CDCs (n=41) and CFs (n=29) at passage 3, in addition to HL-1 cardiomyocytes (n=13). Both CTs and CDCs represent 3 biological replicates. Dim.1 separates the cardiac stromal populations, whereas Dim.2 shows a distinct separation between cardiac stromal and myocyte populations. Gene loadings contributing to each Dim suggest that a small subset of genes explain the cross-group variability captured by Dim.1 and Dim.2. Tcf21 is associated with CTs (emphasised in the blue circle), whereas Tbx5 is associated with CDCs and CFs (emphasised in the brown circle). The separation between cardiac stromal and myocyte populations is reflected by clustering of cardiogenic and cardiomyocyte genes (Nkx2.5, Wif1, Nppa, Myl2 and Myh6), which are emphasised in the pink circle. **(B)** PCA of the three cardiac stromal populations alone: CTs (n=28), CDCs (n=41), CFs (n=27). Both CTs and CDCs represent 3 biological replicates. Gene loadings contributing to each Dim show that Tcf21 is associated with CTs, whereas Tbx5 is associated with CDCs and CFs.

Statistical analysis comparing the three cardiac stromal population revealed that CTs were more similar to CDCs than CFs with CTs showing significantly higher expression of *Vim* (P < 0.05), *Kdr* (P < 0.05), *Mrc1* (P < 0.001), *Tcf21* (P < 0.0001), *Ms4a4d* (P < 0.0001), *Tbx20* (P < 0.0001), *Csf1r* (P < 0.0001), *Mef2c* (P < 0.0001), and *Medag* (P < 0.05), but lower expression of *Tbx5* (P < 0.0001) and *Acta2* (P < 0.01), compared to CDCs. In comparison to the CFs, CTs showed significantly higher expression of *Col1a1* (P < 0.0001), *Pdgfrb* (P < 0.0001), *Hand2* (P < 0.0001), *Pou5f1* (P < 0.05), *Mrc1* (P < 0.001), *Tcf21* (P < 0.0001), *Tbx20* (P < 0.0001), *Ng2* (P < 0.01), *Mef2c* (P < 0.05), *Medag* (P < 0.001), and *Gata4* (P < 0.0001), but lower expression of *Isl1* (P < 0.0001) and *Tbx5* (P < 0.01) (**Figure 5.A**).

**Figure 5.**
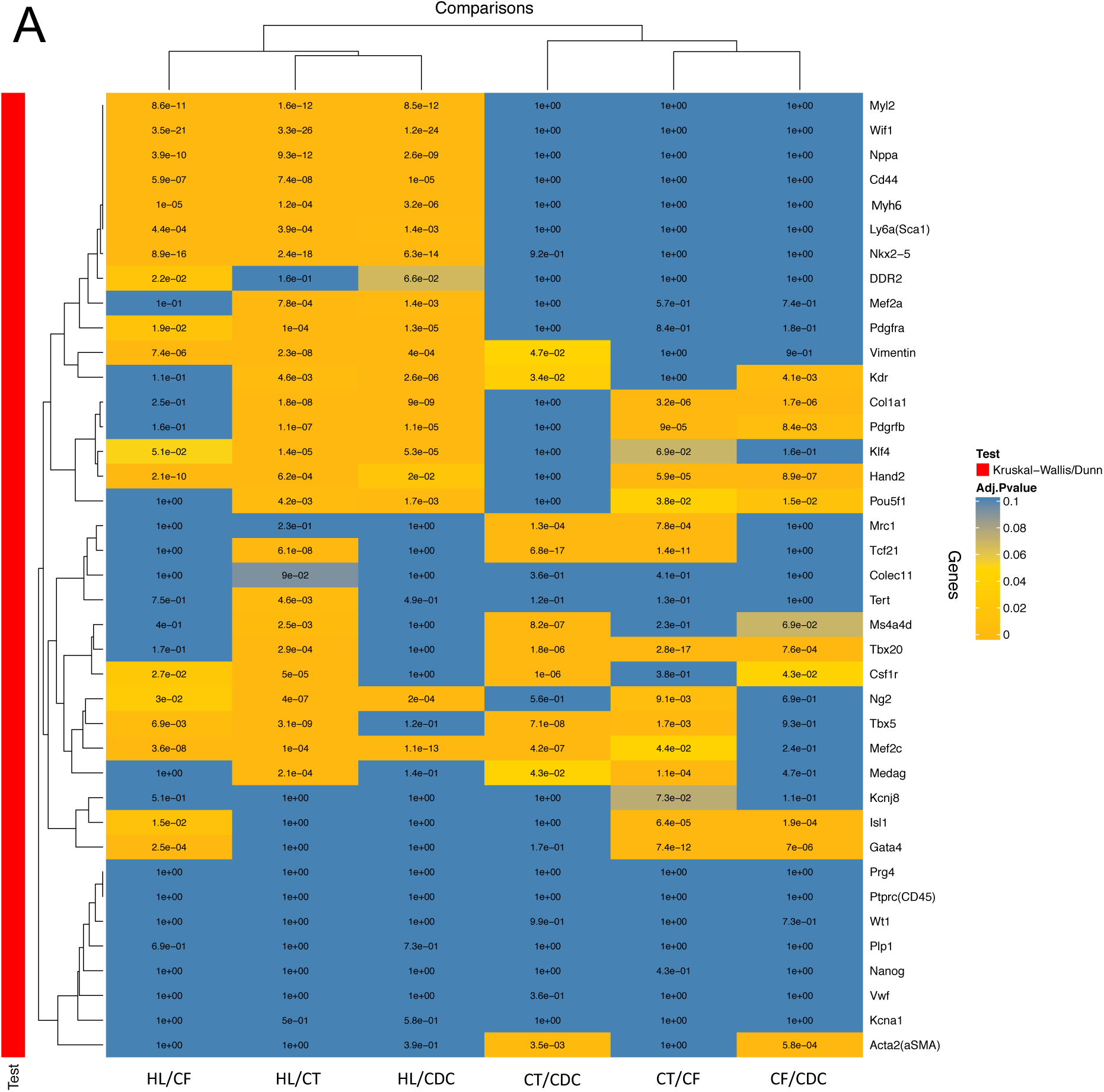
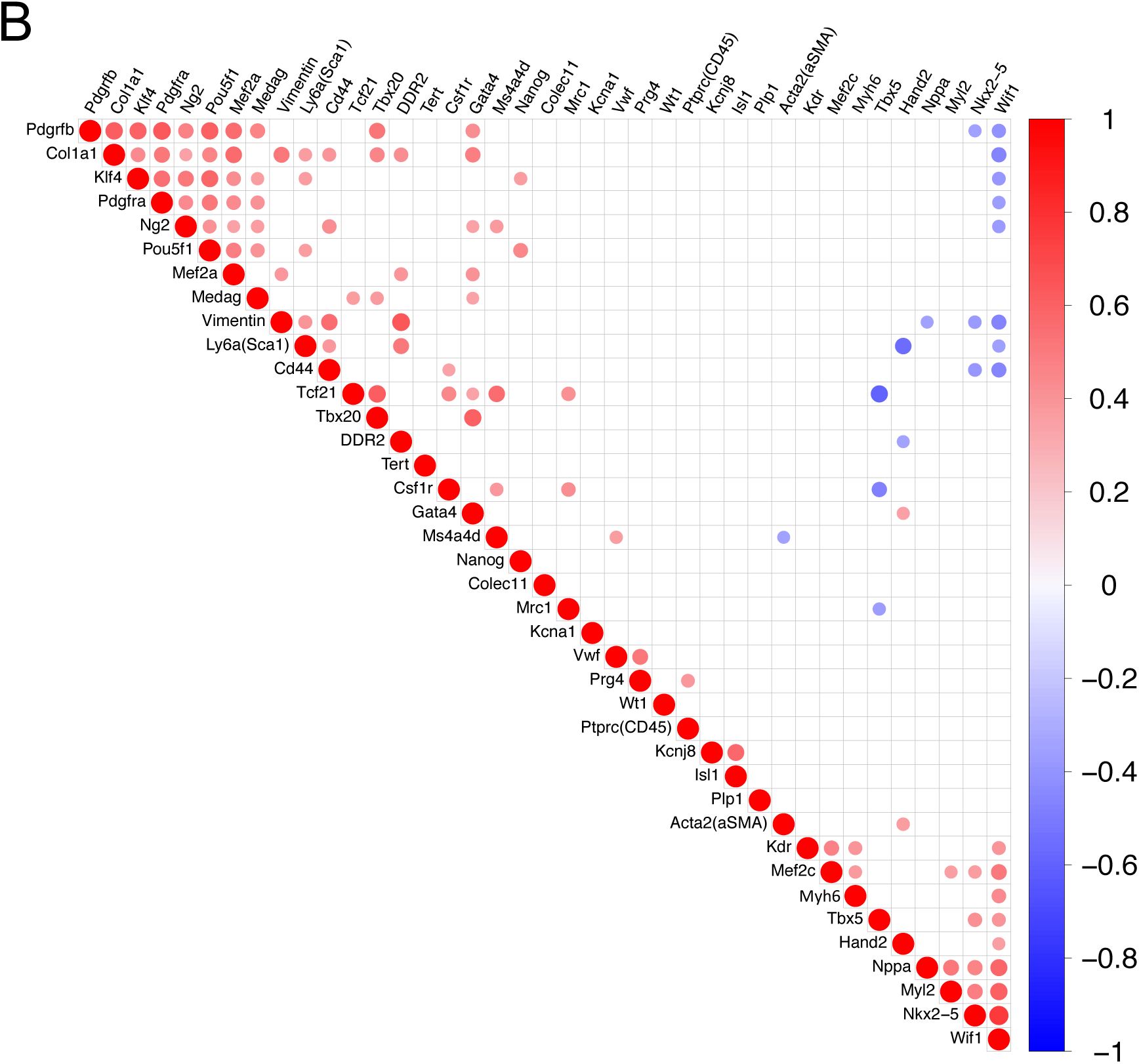
Statistical analysis of the single-cell qRT-PCR and gene correlations. **(A)** Statistical analysis of the four cell populations shown: CTs (n=55), CDCs (n=41) and CFs (n=29) at passage 3, in addition to HL-1 cardiomyocytes (n=13). Both CTs and CDCs represent 3 biological replicates. The heatmap illustrates P values from 0 to 0.1 (orange to blue is low to high). The genes were grouped by a hierarchical clustering algorithm based on the underlying co-expression pattern. **(B)** Gene correlation plot, derived from the single-cell qRT-PCR data, is represented by the colour of the dots (blue is negative; red is positive). Both the diameter and colour intensity of each dot are proportional to the level of correlation.

Finally, we performed co-expression correlation to further support the inferred biological functions of the stromal and cardiomyocyte populations (**Figure 5.B**). Correlation plot analysis showed a polarisation between the expression of some fibroblast-associated markers and cardiogenic transcription factors, and a positive correlation between the expression of cardiogenic transcription factors and cardiomyocyte markers. *Tcf21* expression showed a positive correlation with the expression of *Tbx20*, *Ms4a4d,* and macrophage-associated genes *Csf1r* and *Mrc1*. This is attributed to the high co-expression of these genes in CTs. In addition, expression of pericyte-associated gene *Kcnj8* showed a positive correlation with the expression of *Isl1* due to their co-expression pattern in a small number of CFs. Collectively, these data confirm the co-expression of cardiogenic markers, associated predominantly with CTs and CDCs, as distinct from the gene profile that characterises CFs. Additionally, the enrichment of *Tcf21* expression in CTs further delineates CTs from CFs.

## Discussion

### CTs have reduced expansion time and a superior survival potential following serum starvation

Low survival and retention of transplanted cells following ischaemic events is mainly due to the harsh conditions within the infarct region that include hypoxia, inflammation, and nutrient restriction. Serum in cell culture media provides amino and fatty acids and its withdrawal is therefore, considered a form of nutrient restriction and serum starvation is an effective way of inducing cell apoptosis^30^. Here, we found that all three cardiac stromal populations continued to grow following serum starvation with no significant difference in cell numbers detected at day 3. However, CF cell numbers at day 10 were similar to the cell number seeded initially suggesting substantial cell death. At day 10, CT cell numbers were significantly higher than CFs but not CDCs, suggesting that CTs have a superior survival potential to that of CFs. This is consistent with the single-cell qRT-PCR results showing that CTs and CDCs share a more similar gene expression profile in comparison to the CFs. Tolerance to serum starvation has also been observed in MSCs under both normoxic and hypoxic culture conditions^31,32^.

Reduced robustness was noticed during routine culture of CFs in comparison to CTs and CDCs as shown by increased time to reach confluency and higher numbers of floating dead cells in culture (data not shown). The CFs displayed reduced recovery following single-cell sorting for the single-cell qRT-PCR experiment, in comparison to CTs and CDCs. It is difficult to pinpoint what is responsible for this fragility due to difference in gene expression, cell culture media and coating substrate. Differential effects of various culture coating substrate types on cell proliferation have been shown^33^. However, standardising the culture conditions of all cell types would have been counterproductive as our aim was to compare the populations as they would be cultured in other labs based on their recommended and published culture conditions. Ideally, the survival potential of these different stromal population should be compared following transplantation into infarcted hearts or in *in vitro* conditions that more closely mimic the *in vivo* environment following MI. Overall, our data suggested that CTs and CDCs were more robust than CFs, both in long-term cell expansion and survival under serum starvation. The survival potential of cardiac stromal populations may differ in response to stress *in vitro*, which can be an indication of their *in vivo* behaviour following transplantation. Additionally, the reduced expansion time needed to generate CTs, compared to CDCs, indicated that a CDC-like population for therapeutic use could be achieved more efficiently.

### CPCs show a fibroblast phenotype with no detection of endothelial cells or pericytes

A study by Skelly *et al*. 2018 examined the single-cell transcriptional profile of adult mouse ventricular nonmyocytes to identify distinct cell populations^19^. Subpopulation heterogeneity within the cardiac fibroblast population was described where most fibroblasts “fibroblasts 1” were characterised by high expression of *Medag, Ms4a4d, Pdgfra* and *Tcf21*, while a smaller distinct cluster “fibroblasts 2” expressed higher levels of *Tbx20*, *Wif1* and *Prg4*. Although we found that *Tbx20* was highly expressed in both CTs and CDCs, *Wif1* and *Prg4* showed little to no expression in all stromal populations. Moreover, *Medag* and *Pdgfra* expression was detected in all three populations, whereas *Ms4a4d* and *Tcf21* was only found in CTs and CFs. However, CFs showed less consistent expression of *Ms4a4d* compared to CTs. This suggests that none of the three stromal populations fit the expression pattern of “fibroblasts 2” and that CTs were the best match to the expression pattern of “fibroblasts 1”. However, an important consideration to make is that the study by Skelly *et al*. examined ventricular rather than atrial nonmyocytes and used freshly isolated cells rather than passaged cells which may undergo phenotypic drift *in vitro*.

Additional mesenchymal- and fibroblast-associated genes were assessed including *Vim*, *Ddr2*, *Acta2*, *Cd44*, and *Col1a1*. αSMA, encoded by *Acta2,* is a protein commonly used to identify smooth muscle cells, pericytes and myofibroblasts, which acquire a contractile phenotype similar to that of smooth muscle cells upon differentiation^34^. *Vim* was the only gene expressed in all the screened cells of all populations. Vimentin is an intermediate filament protein that has been widely used as a reliable marker of mesenchymal cells, highly migratory cells derived by EMT, and has a protective role against nuclear rupture and DNA damage during migration^35^. None of these markers showed specificity to a certain population. *Vim*, *Ddr2* and *Acta2* were also expressed in HL-1 cardiomyocytes, an adult immortalised atrial cell line that has been shown to be similar to a mitotic embryonic cardiomyocyte rather than a mature adult cell^36^, as reflected by their immature energy metabolism^37^. The expression of mesenchymal- and fibroblast-associated genes in HL-1 cardiomyocytes may reflect the lack of specificity of these markers in *in vitro* cultures, the immature phenotype of HL-1 cardiomyocytes and/or their dedifferentiation in culture. Genomic instability is a common setback associated with the use of cell lines in research^38^. Nonetheless, the above data suggest a fibroblast-like phenotype of all stromal populations. A combination of *Pdgfra, Col1a1 and Medag,* which were expressed in the stromal cells but not the HL-1 cells, might be more reliable markers for identifying *in vitro* fibroblast cultures.

Commonly used markers to identify pericytes (e.g. *Pdgfrb* and *Ng2*) are relatively non-specific. However, *Kcnj8* and *Colec11* have been identified as genes that show higher expression in pericytes relative to other cell types expressing them^19^. We found little or no expression of *Kcnj8* in any population. *Colec11* showed sporadic expression, whereas *Ng2* and *Pdgfrb* showed high expression in all three cardiac stromal populations. Therefore, there is no conclusive evidence to suggest that these stromal populations are pericytes in their phenotype. Our data also suggest that the specificity of *Kcnj8* and *Colec11* expression to pericytes might be superior to that of commonly used markers. Moreover, the expression of *Kcnj8* was positively correlated with *Isl1*, attributable to co-expression in a small number of CFs. ISL1-mediated differentiation of coronary pericytes to coronary artery smooth muscle cells has been described in the embryonic heart^39^. Although *Isl1* expression is detected in the developing heart and multipotent ISL1^+^ progenitors play an important role in giving rise to cardiac lineages during development^40^, ISL1^+^ cells are very rare in adult human and murine hearts^41,42^.

The three cardiac stromal populations had very little or no expression of markers of cardiomyocytes (*Nppa*, *Myh6*, and *Myl2*), haematopoietic (*Cd45*), or endothelial cells (*Cdh5* and *Vwf*). Expression of *Nppa* and *Myh6* was found in few CDCs (2 *Nppa*^+^ cells, n=41) and CTs (4 *Myh6*^+^ cells, n=55). Given that these cells are at passage 3, it is highly unlikely that the cells expressing *Nppa* or *Myh6* cells are cardiomyocytes that survived the isolation protocol. This expression may be a result of spontaneous differentiation in culture or may reflect some progenitor status. In addition, *Kdr* expression, a marker of endothelial cells, was detected in the CTs, CFs and HL-1 cardiomyocytes. During development, KDR^+^ progenitors, deriving from the MESP1^+^ mesodermal lineage, have a cardiovascular tri-lineage potential^43,44^. It has been shown by lineage tracing that precursors expressing *Gata5* and *Wt1* give rise to the adult SCA1^+^ SP^+^ population (PDGFRα^+^/CD31^-^), whereas the adult SCA1^+^ SP^-^ population (PDGFRα^-^/CD31^+^) is mainly derived from precursors expressing *Kdr^9^.* However, CTs lack the expression of other endothelial markers as observed in the SCA1^+^ SP^-^ population. *Kdr* expression has also been described in coronary adventitial progenitors with cardiogenic potential^45^ and in endothelial progenitors^46^. Therefore, *Kdr* expression may reflect some progenitor status as opposed to the presence of endothelial cells. CDCs have been examined at the single cell transcriptomics level^47^ and characterised as mesenchymal/stromal/fibroblast-like, with a small subset of endothelial-like cells. Notably, transplantation of SCA1+ CDCs but not SCA1− CDCs enhanced cardiac function after MI, highlighting the previously unappreciated functional differences between CDC subpopulations.

A few cells co-expressing *Wt1* and *Tcf21* were detected in both the CF and CT populations. In the developing heart, cells of the epicardium are identified through the expression of *Wt1*, *Tbx18*, *Tcf21*, and *Raldh2*^48^. It has been demonstrated that the majority of TCF21^+^ epicardium-derived cells (EPDCs) are committed to the cardiac fibroblast lineage and that *Tcf21* expression persists in cardiac fibroblasts of the adult heart^49–51^. Although WT1 is downregulated following gestation^52,53^, its expression is still detected in the epicardium and a subset of coronary endothelial cells in the healthy adult heart^50,54^. Enrichment of the epicardial markers *Wt1* and *Tcf21* has been shown in adult cardiac fibroblasts^17^. More recent studies showed that *Wt1* is expressed in more cell types than previously appreciated with a study describing a Wt1b^+^ macrophage subpopulation that plays an essential role in zebrafish heart and fin regeneration^55^. In our data, *Wt1* expression did not show a correlation with *Mrc1* or *Kdr* but all cells expressing *Wt1* were also positive for *Tcf21* suggesting a fibroblast phenotype.

Expression of stem cell-related markers (*Pou5f1*, *Nanog*, *Klf4*, and *Tert*) was found in all three cardiac stromal populations. In addition, *Pou5f1*, *Nanog*, and *Klf4*, but not *Tert*, also showed expression in HL-1 cardiomyocytes. The expression of these stem cell-associated genes (*Pou5f1*, *Klf4*, *Nanog*) has been observed in mouse cardiac stromal populations and rat CDCs^56^. A sporadic expression of these markers has also been noted in primary neonatal cardiomyocytes^9^. Other studies have also reported the expression of *Pou5f1*^57^*, Klf4*^58^*, Nanog*^59,60^ in adult differentiated cells. Furthermore, *Tert* is activated in human and mouse primary fibroblasts to protect against malignant transformation^61^. As stem cell-associated genes have been implicated in tumorigenesis and proliferation^62,63^, it is not surprising to observe their expression in the HL-1 cell line or the stromal populations.

### *Tcf21* versus *Tbx5* polarity may represent functional heterogeneity in fibroblast populations and reveal a macrophage phenotype in a CT subpopulation

Noseda *et al*.^9^ revealed an enrichment of *Tcf21* and *Pdgfra*, the cardiogenic genes *Gata4, Mef2a/c, Tbx5/20 and Hand2*, and a lack of *Cdh5* and *Kdr* in the SCA1^+^ SP^+^ population in comparison to total SCA1^+^ and SCA1^+^ SP^-^ populations. In our study, a lack of *Cdh5* expression was observed in all cardiac stromal populations. Both CTs and CDCs showed high expression of the cardiogenic transcription factors *Gata4*, *Tbx20*, *Hand2*, and *Mef2a* with CTs expressing *Mef2c* and CDCs expressing *Tbx5*. Interestingly, among these transcription factors, *Tbx5* showed the lowest expression in the SCA1^+^ SP^+^ population. Moreover, these results are consistent with the observations of Furtado *et al*. showing an enrichment of cardiogenic genes in cardiac fibroblasts^17^. In contrast, CFs had less uniform expression of *Gata4*, *Tbx20*, and *Hand2*. The PCA analysis showed an interesting polarisation between *Tcf21* and *Tbx5*, which was also reflected on the correlation plot showing a strong negative correlation between the two genes. Indeed, the separation in clustering of CTs and CFs can be attributed to the expression of these two genes. Although it is unclear what this polarisation means, it is an interesting observation that warrants further investigations in future studies as it may reflect potential functional heterogeneity.

In addition to the enrichment of *Tcf21* in CTs as opposed to other stromal populations, the macrophage-associated gene *Mrc1* was detected in a subset of CTs. This was also reflected by the positive correlation between *Mrc1* and *Tcf21* versus the negative correlation between *Mrc1* and *Tbx5*. *Mrc1* has been reported to show higher expression in macrophages, relative to other cardiac cell types^19^. *Csf1r*, enriched in M2-polarised macrophages^64^, was expressed in all stromal populations, albeit at significantly higher levels in CTs. This suggests the possibility of a macrophage-like subpopulation in CT populations, and potentially in CDC and CF populations as well. *Csf1r* expression has also been used as an indicator of embryonic myeloid lineage commitment^65^. However, cells expressing *Mrc1* or *Csf1r* showed co-expression of fibroblast markers and did not show clear separation in clustering. Cells expressing *Mrc1* or *Csf1r* may therefore represent the transdifferentiation of macrophages in the stromal isolations into fibroblasts or *vice versa*. Indeed, the transdifferentiation of macrophages to fibroblast cells has been described *in vivo*^66^. Moreover, hybrid mesenchymal phenotypes have been previously described where fibroblast-associated genes are expressed in minor subpopulations of macrophage or endothelial cells^19^. However, due to the lack of *Cd45* expression in CTs, it is more likely that macrophage-associated genes were upregulated in a subset of fibroblasts rather than *vice versa*. This phenomenon has been previously reported in *in vitro* fibroblast cultures^67^.

Our results suggest that (1) CTs were similar to CDCs and differed from CFs in their expression profile, enrichment of cardiogenic factors and robustness; (2) CTs, CDCs, and CFs resembled “fibroblasts 1”, which are the predominant fibroblast subtype in the adult murine heart; (3) CDCs clustered between CTs and CFs which separated based on their expression of *Tcf21* and *Tbx5*, respectively; (4) CTs were enriched for the macrophage-associated genes *Mrc1* and *Csf1r*, which may be attributed to transdifferentiation of macrophages to or from a fibroblast phenotype in a subset of CTs; (5) PCA analysis revealed a transcriptional gradient suggesting that heterogeneity exists within fibroblast subtypes which may reflect functional heterogeneity. Overall, our findings highlight the heterogeneity of the stromal compartment of the heart and how the isolation protocol may enrich for specific cell subpopulations, which may ultimately affect the therapeutic potential of the cells. Furthermore, the decreased culture time required to generate CTs, as compared to that for the CDCs, suggests that a CPC population for therapeutic use could be achieved more efficiently.

## Methods

### 1. Mice

C57BL/6 mice (Harlan, Oxon, UK) were kept under in a 12-hour light-dark cycle, and controlled conditions of temperature, humidity, with free access to water and chow. All animal procedures were reviewed and approved by the University of Oxford Animal Welfare and Ethical Review Board and conforms to the Animals (Scientific Procedures) Act 1986 incorporating Directive 2010/63/EU of the European Parliament.

### 2. Isolation and expansion of mouse CPCs

Mice were terminally anesthetised with isoflurane and hearts were isolated and washed with Dulbecco’s phosphate buffered saline (DPBS) containing 50mg of primocin (antimicrobial agent; InvivoGen). Mouse atrial appendages were dissected and minced mechanically into 1 - 2 mm^3^ pieces. For CT isolation, the tissue pieces were then transferred into a digestion mix (0.1% trypsin and 0.1% Collagenase II (Calbiochem, 286U/mg) in DPBS) and incubated in a water bath at 37°C for a total of 40 minutes. Every 10 minutes the digestion mix was mechanically triturated by pipetting, left on ice for 1 minute to settle and the supernatant was collected. Fresh digestion mix was added again to the tissue pieces for a total of 4 digestions. At the final digestion step, sterile 19G needles and 1ml syringes were used to triturate the tissue. After each digestion, the supernatant was neutralised, and the cell suspension was resuspended in fresh complete explant medium (CEM; Iscove’s modified Dulbecco’s medium supplemented with 20% FBS, 100U/ml penicillin, 100μg/ml streptomycin and 2mM L-glutamine, ThermoFisher)^11^ to be plated in a 12-well plate through a 40µm cell strainer. The wells were pre-coated with 50µl per cm^2^ of fibronectin (2µg/ml in DPBS; Sigma-Aldrich) and cells were allowed to attach for 3 days without media changes. For CDC isolation^11^, the tissue pieces were digested in 0.05% Trypsin-EDTA (ThermoFisher) for 4 minutes at 37°C then neutralised with CEM. Processed tissue pieces were plated on fibronectin-coated (3μg/ml in DPBS) 6-well plates containing CEM and explants were cultured for 25-30 days at 37°C in 5% CO2 with media changes every 2-3 days. When explant-derived cells (EDCs) reached 70-80% confluency, they were harvested by washing with 0.53mM Versene (ThermoFisher) before a trypsin digestion. EDCs were resuspended in Cardiosphere Growth Medium (CGM), comprising 65% Dulbecco’s modified eagle medium (DMEM/F12), 35% IMDM, 7% foetal bovine serum (FBS; ThermoFisher), 2% B27 (ThermoFisher), 25ng/ml cardiotrophin (Peprotech EC), 10ng/ml epidermal growth factor (EGF; Peprotech EC), 20ng/ml basic fibroblast growth factor (FGF; Promega) and 5 units thrombin (Sigma-Aldrich), at a concentration of 2x10^4^ cells/ml and seeded as 25µl droplets on an un-coated lid of a Petri dish. DPBS was added to the Petri dishes to keep the hanging drops humid throughout 3 days of culture allowing the spherical multicellular clusters, known as cardiospheres, to form. Cardiospheres were collected by elution and gentle pipetting with DPBS and were plated in fibronectin-coated 12-well plates with CEM allowing them to spontaneously release CDCs.

### 3. Cell culture

Primary CPCs were cultured in CEM and plated on fibronectin-coated flasks. HL-1 cardiomyocytes^36^ were maintained in Claycomb medium (Sigma-Aldrich), supplemented with 100U/ml penicillin, 100μg/ml streptomycin and 2mM L-glutamine (ThermoFisher), 100µM norepinephrine (Sigma-Aldrich) in 30mM L-ascorbic acid (Sigma-Aldrich), and 10% FBS and plated on flaks pre-coated with 0.02% (wt/vol) gelatin (Sigma-Aldrich) containing 5µg/ml fibronectin. C57BL/6 adult mouse atrial primary cardiac fibroblasts, obtained from Cell Biologics (USA), were maintained in complete fibroblast medium (Cell Biologics – USA) supplemented with the provided supplement kit: FGF, hydrocortisone, L-glutamine, Antibiotic-Antimycotic Solution, and 10% FBS and plated on flasks pre-coated with 0.02% (wt/vol) gelatin. For serum starvation experiments, cells were switched to serum-free media for 72 hours.

### 4. Single-cell qRT-PCR

Single cells were sorted directly (FACSAria II) into 96-well plates containing the reaction mixture for pre-amplification, using CellsDirect One-Step qRT–PCR Kits (ThermoFisher). Pre-amplification was performed in a Veriti Thermal Cycler (Applied Biosystems) for 22 cycles. As negative controls, at least 3–5 non-template samples were included in each run at the pre-amplification stage. Quantitative amplification was performed using Dynamic Array chips for 48 assays x 48 samples, the BioMark HD system (Fluidigm) and TaqMan probes (Appendix D) according to manufacturer’s instructions. Each sample was normalised to ΔCt using the expression of *Ubc*. As described in Noseda *et al*. 2015^9^, data were plotted in colour-coded heatmaps of inverted ΔCt values (blue is low or absent; red is high). Samples were ordered based on cell type, and the genes grouped by a hierarchical clustering algorithm based on the underlying co-expression pattern. Differences between samples were investigated using PCA. PCA applies multiple linear transformations (singular value decomposition) to the expression profiles (standardised ΔCt values) of individual samples, and identifies a series of PCs that elucidate the most distinguishing features between the samples. The linear projections (PC scores) attempt to maximise the variation between the samples, whereas the coefficients of those projections (PC loadings) measure the importance of genes in defining the underlying variability associated with each component.

#### Taqman probes used for single-cell qRT-PCR

|  |  |  |  |
| --- | --- | --- | --- |
| *Mesp1* | Mm00801883_g1 | *Nkx2-5* | Mm00657783_m1 |
| *Mef2c* | Mm01340842_m1 | *Myh11* | Mm00443013_m1 |
| *Mef2a* | Mm01318990_m1 | *Cnn1* | Mm00487032_m1 |
| *Hand1* | Mm00433931_m1 | *αSMA (Acta2)* | Mm01546133_m1 |
| *Islet1 (Isl1)* | Mm00627860_m1 | *Vim* | Mm01333430_m1 |
| *Tbx20* | Mm00451515_m1 | *Cd44* | Mm01277163_m1 |
| *Gata4* | Mm00484689_m1 | *Pdgfrb* | Mm00435546_m1 |
| *Hand2* | Mm00439247_m1 | *Ng2* | Mm01283063_m1 |
| *Tbx5* | Mm00803518_m1 | *Ddr2* | Mm00445615_m1 |
| *Ly6a (Sca1)* | Mm00726565_s1 | *Medag* | Mm00551008_m1 |
| *Wt1* | Mm00460570_m1 | *Ms4a4d* | Mm00656404_m1 |
| *Klf4* | Mm00516104_m1 | *Col1a1* | Mm00801666_g1 |
| *Pou5f1 (Oct4)* | Mm03053917_g1 | *Prg4* | Mm01284582_m1 |
| *Nanog* | Mm02019550_s1 | *Wif1* | Mm00442355_m1 |
| *Tert* | Mm0352136_m1 | *Colec11* | Mm01289834_m1 |
| *Kdr* | Mm00440111_m1 | *Kcnj8* | Mm00434620_m1 |
| *Cdh5* | Mm00486938_m1 | *Mrc1* | Mm01329362_m1 |
| *Vwf* | Mm00550376_m1 | *Cx3cr1* | Mm00438354_m1 |
| *Pdgfra* | Mm01211685_m1 | *Csf1 r* | Mm01266652_m1 |
| *Tcf21* | Mm00448961_m1 | *Kcna1* | Mm00439977_s1 |
| *Myl2* | Mm00440384_m1 | *Plp1* | Mm01297210_m1 |
| *Myh6* | Mm00440359_m1 | *Ptprc (Cd45)* | Mm01293577_m1 |
| *Nppa* | Mm01255748_g1 | *Ubc* | Mm01201237_m1 |
| *Hmbs* | Mm01143545_m1 |  |  |

**Supplementary Table 1.**
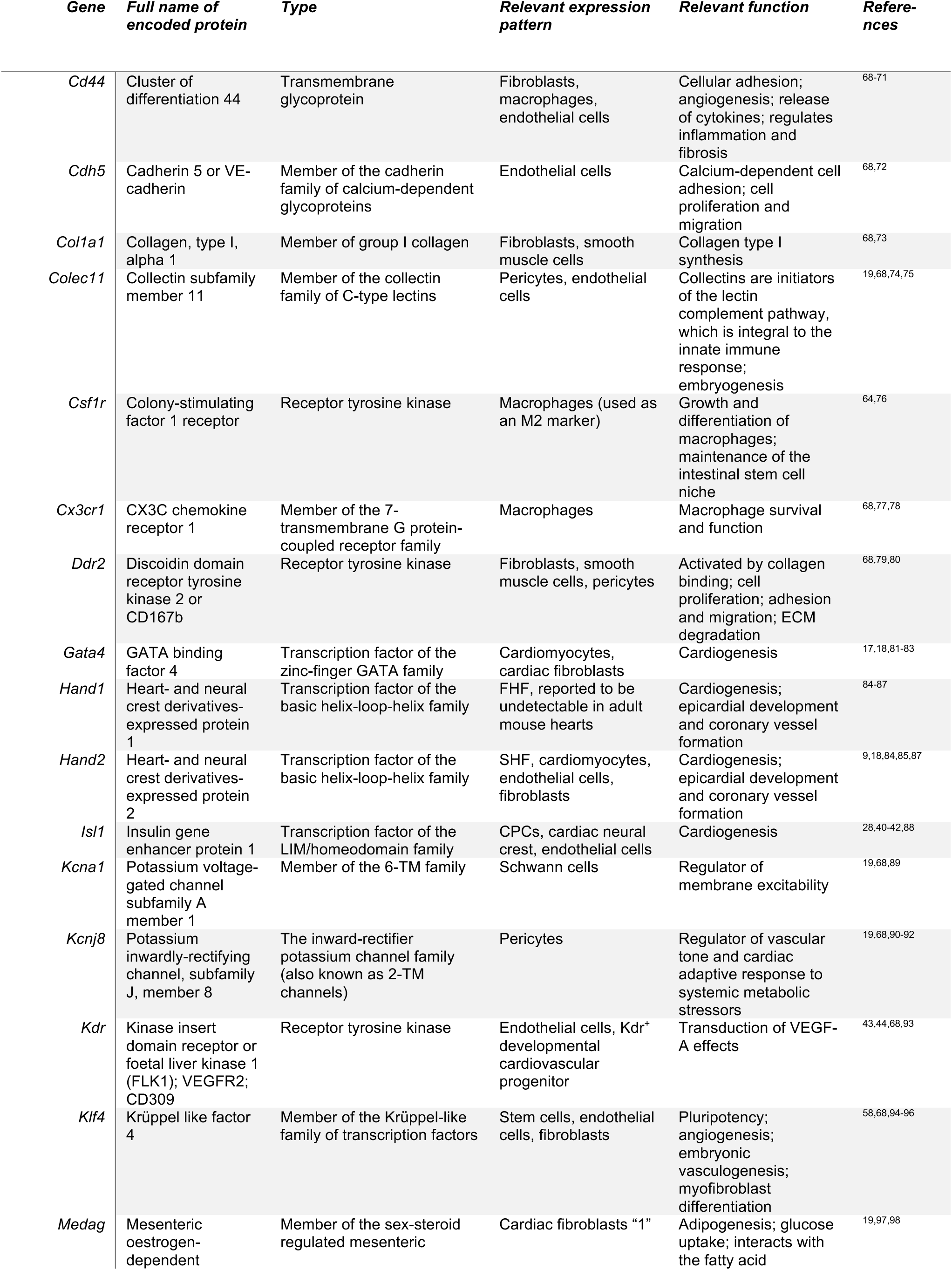
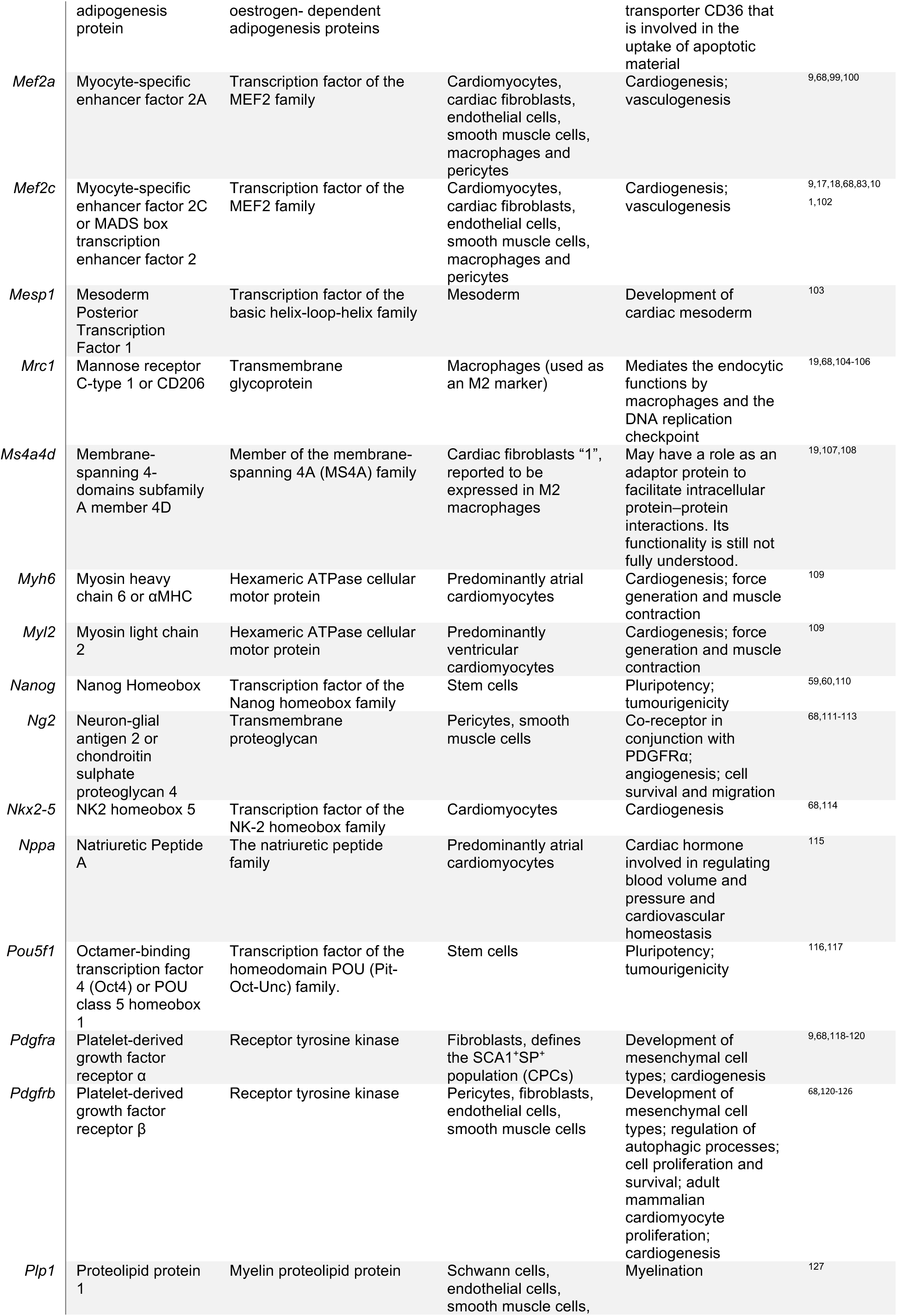
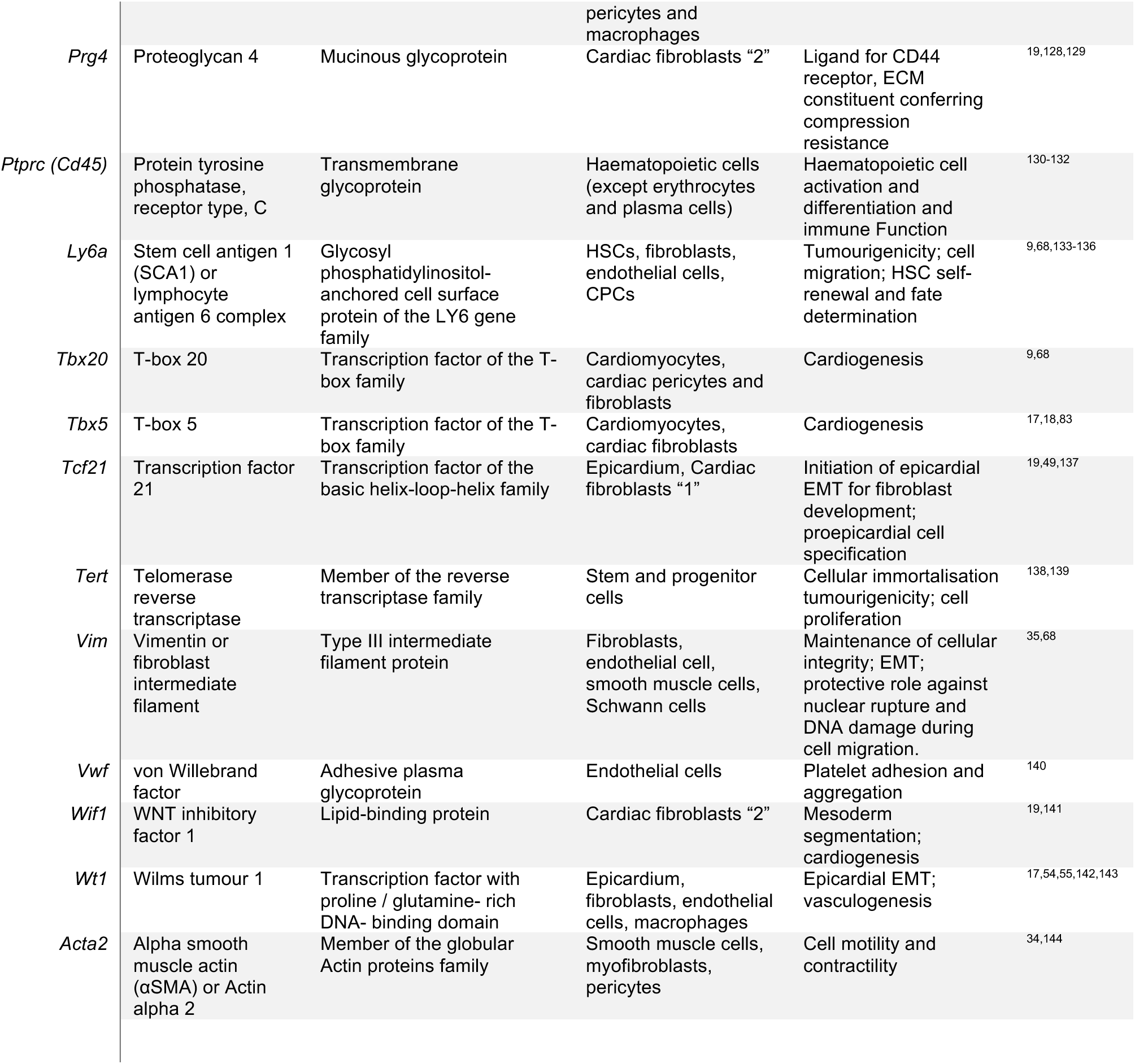
Description of the markers used in the single-cell qRT-PCR characterisation.

## Author contributions

Conceptualisation: RA, NS, CC. Investigation: RA, PCG, SS, MN. Funding acquisition: RA. Writing: RA, CC.

## Acknowledgements

We would like to thank Dr Andrea Massaia for the R script used in the single-cell qRT-PCR analysis. This work was supported by the King Faisal Specialist Hospital & Research Centre (PhD scholarship to RA).

